# Age-dependent genetic architecture underlines similar heritability of body size in sticklebacks

**DOI:** 10.1101/2021.04.19.440387

**Authors:** Antoine Fraimout, Zitong Li, Mikko J. Sillanpää, Juha Merilä

## Abstract

Heritable variation in traits under natural selection is a prerequisite for evolutionary response. While it is recognised that trait heritability may vary spatially and temporally depending under which environmental conditions traits are expressed, less is known about the possibility that genetic variance contributing to the expected selection response in a given trait may vary at different stages of ontogeny. Specifically, whether different loci underlie the expression of a trait throughout development – thus providing an additional source of variation for selection to act on – is unclear. Here we show that the heritability (*h^2^*) of body size, an important life history trait, remains constant across ontogeny in a stickleback fish. Nevertheless, both analyses of quantitative trait loci (QTL) and genetic correlations across ages show that different chromosomes/loci contribute to this heritability in different ontogenic time-points. This suggests that body size can respond to selection at different stages of ontogeny but that this response is determined by different loci at different points of development. Hence, this illustrates the notion that diverse genetic architectures may underline similar (expected) phenotypic outcomes, and that similar selection pressures may lead to genetically heterogeneous responses depending on what life stage selection is acting on.

## Introduction

How predictable is evolution? This question has intrigued evolutionary biologists for over a century [1–4] and although much progress has been made towards answering it, more remains to be learned about conditions and processes influencing the degree of predictability in evolutionary responses [5–8]. Understanding when and why evolution is predictable is not only of pure academic interest, but also of practical utility. For instance, when dealing with the most pressing environmental problem of our times, the global change, there is a need to predict and understand if and how different species’ and populations will be able to adapt to changing environmental conditions [9].

Both empirical and theoretical work have established that when populations repeatedly and independently adapt to similar environmental conditions from the *same* pool of ancestral genetic variation, evolution is often (but not always) predictable: parallel adaptations evolve both at phenotypic and genetic levels [10]. A prime example of this is the loss lateral armour plates in the three-spined stickleback (*Gasterosteus aculeatus*) in response to colonization of freshwater habitats from marine environments; this adaptation is predictably underlined by parallel genetic changes in the Ectodysplasin (EDA) gene [11–13]. Inspired by this success story, the three-spined stickleback has become ‘the’ empirical model system to study predictability of evolution (*e.g*. [14–17]). However, when populations repeatedly and independently adapt to similar environmental conditions from *different* pools of ancestral genetic variation, evolution becomes much less predictable (*e.g*. [8, 17]). In fact, theory suggests that the predictability of evolution - defined as the probability of parallel evolution from standing genetic variation - is a positive function of effective population size (*N_e_*) in the ancestral population [18, 19] which, in turn, is a positive function of gene flow [20–22]. Therefore, heterogeneity in pools of standing genetic variation can be expected to lower the likelihood of genetic parallelism, as evidenced also by empirical observations (*e.g*. [8]).

While a decade of evolutionary genetics research has explored factors influencing the probability of parallel evolution (*e.g*. [1, 3, 7, 10, 16, 23–26]), and even demonstrated that phenotypic similarity can be underlined by different genes (*e.g*. [8, 27–31]), less attention has been paid on age-dependent heterogeneity of genetic architecture as a potential source of nonparallel evolution at the genetic level. Indeed, if different genes contribute to the expression of a given trait at different ontogenetic stages and natural selection acts on the trait at different stages of ontogeny in different locally adapted populations, the expected outcome of this would be among population divergence in genetic architectures. Hence, under these premises, similar selection pressures on the trait in different populations would be expected to promote genetic nonparallelism, that is, reduce the predictability of evolutionary outcomes at the genetic level.

The aim of this study was to explore if selection acting on a phenotypic trait (body size) in different times of ontogeny would lead to similar expected phenotypic responses to selection, and whether these responses would be underlined by similar or different genetic architectures. To this end, we analysed heritability and the contributions of different chromosomes to body size variation over ontogeny in three F_2_-generation inter-population crosses of nine-spined sticklebacks (*Pungitius pungitius*). Significant additive genetic variance and approximately constant heritability of body size across ages would suggest that this trait could respond to directional selection at different stages of the ontogeny. If the same chromosomes contribute to body size variation at different ontogenetic stages and in all different crosses, this would suggest that evolutionary outcome of selection on body size would be predictable irrespectively of population and ontogenic stage the selection is acting on. If on the other hand, different chromosomes underline variation in the body size at different ontogenetic stages and crosses, this would suggest that outcome of similar selection pressure in respect to ontogeny and population would be unpredictable at genetic level.

## Material and Methods

### Study species, crosses & phenotypic data

The nine-spined stickleback is a teleost fish with a wide distribution range across the northern hemisphere [32]. In Europe, ancestral marine *P. pungitius* populations have colonized multiple freshwater ponds and lakes since the end of the last glaciation ca. 11,000 years ago [33–35]. Isolated pond populations of *P. pungitius* have repeatedly evolved extreme morphological [8, 36] and behavioural [37] phenotypes, making them particularly interesting for the study of parallel adaptive evolution [8, 17, 38]. Marine and freshwater *P. pungitius* display also contrasting growth trajectories: while the former ecotype shows fast growth to an early maturation at small final size, the latter exhibits prolonged growth and delayed maturation at larger size [39].

Here, we analysed growth trajectories in three independent F_2_ inter-population crosses used also in previous studies exploring the genetic architecture of various quantitative traits in *P. pungitius* [8, 40–42]. Briefly, grandparental (F_0_) individuals were collected from the wild: three different females from a marine population (Helsinki, Finland; 60°13’N, 25°11’E) and three males from three different freshwater ponds: Rytilampi (Finland, 66°23’N, 29°19’E), Pyöreälampi (Finland, 66°15’N, 29°26’E) and Bynästjärnen (Sweden, 64°27’N, 19°26’E). Wild grandparents were mated artificially in the laboratory and F_1_-offspring were reared in 1.4-l tanks in an Allentown Zebrafish Rack Systems (Aquaneering Inc., San Diego, USA; see detailed rearing conditions in [42]). F_2_ offspring were produced by mating single randomly chosen mature male and female in each cross (see detailed rearing conditions in [42]) and in total, the Helsinki x Rytilampi cross (HEL x RYT; mating: July-Sep 2006, F_2_ breeding: Sept-Oct 2008) consisted of 274 F_2_-offspring, the Helsinki x Pyöreälampi (HEL x PYÖ; mating: July-Sep 2012, F_2_ breeding: July 2012-Apr 2013) 278 F_2_-offspring and the Helsinki x Bynästjärnen (HEL x BYN; mating: Nov 2013-Jan 2014, F_2_ breeding: Nov 2013-Aug 2014) 307 F_2_-offspring.

Growth data for all individuals was obtained by measuring the body size of each fish at different time points throughout development. Individuals from HEL x PYÖ and HEL x BYN crosses were measured at nine time points at 4, 8, 12, 16, 20, 24, 28, 32 and 34 weeks post-hatching. Fish from the HEL x RYT cross were measured at seven different time points 2, 6, 10, 14, 18, 22 and 26 weeks post-hatching. Body size was measured as the distance between the tip of the snout and the base of the posterior end of the hypural plate for all individuals from digital photographs including a millimetre scale and using the tps.Dig software [43].

### SNP genotyping

We used restriction-site associated DNA sequencing approach (RADseq; [44]) to obtain a large panel single nucleotide polymorphisms (SNPs) as described in [45]. Two of the crosses used here, HEL x RYT and HEL x PYÖ; correspond to their “HR” and “HP” crosses, respectively. The same methodology as used in [45] was applied to the third (HEL x BYN) cross. In short, phenol-chloroform method [46] was used to extract DNA from alcohol-preserved fin clips to be used for library constructions and fragmented to 300-500bp fragments using the *Pst1* restriction enzyme. DNA fragments were gel purified, ligated to adaptors and library-specific barcodes, after which the individual libraries were pooled for sequencing. Sequencing was conducted using 45-bp single-end strategy on the Illumina HighSeq 2000 platform at BGI Genomics (Hong-Kong Co. Ltd.). Adapters and barcodes were removed from raw reads and the read quality was checked using FastQC [47]. Genotype data (raw) consisted of 21 832, 22 603 and 21 747 SNPs for the HEL x RYT, HEL x PYÖ and HEL x BYN crosses, respectively.

### Phenotypic variation in growth trajectories

We first investigated patterns of growth within each cross to i) determine whether individuals had reached their adult size, and ii) to describe the growth trajectories explaining phenotypic variation in our datasets. To this end, we first summarised growth trajectories using a von Bertalanffy growth curve model [48] as follows:

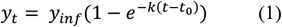

where *y_t_* is the length of an individual at age *t*, *k* the intrinsic growth rate, *t_0_* is the estimated length at age *t* = 0, and *y_inf_* is the asymptotic length, corresponding to the estimated final body size for each cross. Second, we applied an Infinite Dimensional Model (IDM; [49, 50]) to our data using the *InfDim* R package. We fitted the model using the full phenotypic covariance matrix of age-specific body sizes for each cross and estimated the 95% confidence intervals (CIs) via bootstrapping using the *IDM.bootCI()* function.

### Quantitative genetic analyses: genetic variance and heritability

Analytically, longitudinal data such as growth measurements can be viewed either as a character taking on a different value at each discrete age (‘character state trait’), or time-dependent observations describing a continuously varying trajectory (‘function-valued trait’; [49]). Here we used both approaches to analyse the data under a Bayesian framework.

First, we partitioned the phenotypic variance (*V_P_*) of age-specific body size into its additive genetic component (*V_A_*) by fitting an animal model of the form:

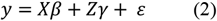

where *y* is the vector of phenotypic values for age-specific body size, *β* is the vector of fixed effect, γ is the vector of random effects, ε is the vector of residual errors and *X* and *Z* the design matrices relating to the fixed and random effects, respectively. In order to improve the accuracy of variance component estimation, we appended the Genomic Relationship Matrix (GRM) constructed from SNP markers to the vector of random effect in our animal models [42]. The GRM was constructed following VanRaden [51] as:

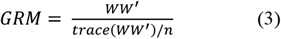

where *W* is the SNP marker matrix of additive coefficients and *n* corresponds to the number of individuals. We used the R package *snpReady* [52] for each dataset to construct the GRM from autosomal loci. After filtering out markers with a minimum allele frequency (MAF) lower than 0.01 and with a maximum of 10% missing data, each GRM was constructed using 12679, 16869 and 15216 informative SNPs for the HEL x RYT, HEL x PYÖ and HEL x BYN crosses, respectively. Each animal model was fitted using the *MCMCglmm* R package [53] using non-informative priors for 1 030 000 Monte Carlo Markov Chain (MCMC) iterations with a burn-in period of 30 000, thinning every 1000th iteration and by adding sex of the individuals as a fixed effect. We fitted univariate animal models for each size at each age separately in all crosses and calculated heritability of age-specific body size from the posterior distribution of the 1000 MCMC samples as:

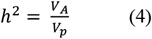

and computed the respective 95% highest posterior density (HPD) intervals using the *HPDinterval()* function in *MCMCglmm*. To allow for meaningful comparison of the magnitude of additive genetic variance across ages, we calculated the coefficients of additive genetic variation (*CV_A_*; [54]):

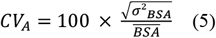

where *σ^2^_BSA_* is the additive genetic variance for body size at age (BSA) and the denominator 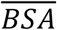 is the phenotypic mean of the body size at a specific age. We applied a penalized spline smoothing function to the full set of MCMC posterior of additive genetic variance and heritability, to account for the dependency of the variance estimates and to confirm the results from the univariate animal models (see *Supplementary material*).

We then re-estimated *V_A_* and *h^2^* following the ‘function-valued trait’ approach by using the method described in [55] implemented in the *dynGP* package (https://github.com/aarjas/dynBGP). Contrary to the univariate animal model approach, this method combines all measurements of body size over time and models the dependency among those measurements using a Bayesian Gaussian process. As such, the method allows to model dynamic variance components and heritability for longitudinal data using SNP data, and to provide reliable uncertainty estimates around the estimated quantities. In these analyses, we followed the methodology of Arjas *et al*. [55] and estimated the GRMs using the *rrBLUP* R package [56] and ran the models for 1 000 000 MCMC-iterations. To account for the effect of sex, we used vectors of residuals from linear regressions of age-specific body size on sex as response variables.

### Quantitative genetic analyses: genetic correlations and G matrix

To estimate genetic correlations across different ontogenetic time-points, we analysed body size across all ages by fitting a multivariate version of the animal model described in equation [2]. Because our growth data correspond to repeated measurements of the same individuals over time, we added a permanent environment effect term in the animal model as a random effect to represent the dependent part of the residuals (caused by those repeated measurements). From the multivariate models, genetic correlations among body size across ontogeny were calculated as:

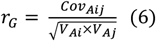

where *Cov_Aij_* is the additive genetic covariance between traits *i* and *j* and *V_Ai_* and *V_Aj_* the additive genetic variances for traits *i* and *j*, respectively. Finally, to investigate the major patterns of genetic variation in growth trajectories a principal component analysis (PCA) of the genetic (co)variance matrix **G** estimated from the multivariate animal models for each cross was performed. Specifically, we were interested in estimating the percentage of additive genetic variance explained by the major axis of genetic variation **g**_max_ and in evaluating the age-specific loadings of **g**_max_. We applied the PCA to the mean **G** matrix of each cross estimated from the multivariate animal models and computed the total variance explained by the first principal component (PC1 = **g**_max_). This also allowed us to estimate the age-specific loadings associated with **g**_max_ and to determine whether ontogeny of body size is integrated (loadings having equal signs) or modular (loadings having variable signs; [57, 58]). Due to the computational burden of MCMC sampling in multivariate animal models using GRM, we reduced the number of observations in HEL x PYÖ and HEL x BYN from nine to seven by removing measurements from age at Week 24 and Week 32. For the same reason, models were run for 203 000 MCMC iterations with a burn-in period of 3000 and thinning every 100th iteration. To confirm the results from univariate models, additive genetic variance and heritability of age-specific body sizes were also computed directly from the multivariate models.

### QTL mapping

We used a single-locus mapping approach [59, 60] to identify the genomic regions underlying variation in body size at each ontogenetic time point. The total effect of each SNP on the phenotype of individuals can be obtained from the regression coefficient of the following Gaussian regression model:

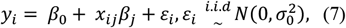

where *y_i_* is the vector of phenotypes (*viz*. age-specific body size) for individuals *i; β_0_* is the phenotypic mean; *x_ij_* is the genotypic value of individual *i* and marker *j* coded as −1, 0 and 1 for the three genotypes AA, AB and BB, respectively; *β_j_* is the effect of the SNP *j*, and ε_i_ is the residual error assumed to be independent and identically distributed (*i.i.d.*) under a normal distribution with zero mean and unknown variance 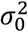. For the case of an inter-cross F_2_ design such as ours, up to four possible alleles can segregate: two alleles A1 and A2 from the dam, and two alleles B1 and B2 from the sire [61, 62]. Consequently, the model described in equation (7) can be stated as:

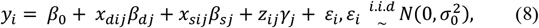

where the substitution effects *β_dj_* and *β_sj_* correspond to the alleles A1 and A2 and B1 and B2 at locus *j* for the dam *d* and sire *s*, respectively, and where *γ_j_* is the dominance effect. In model (8), the matrix of genotype coding system for *x_dij_*, *x_sij_* and *z_ij_* [62] is specified as:

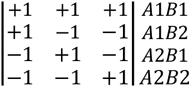

In order to separate the dam and sire alleles – and to obtain additional information of the origin of the QTL effects – parental phasing information must be incorporated in model (8). To this end, we used the data produced in [8] for the same three F_2_ crosses. Briefly, parental and granparental phase were obtained from dense SNP data (see detailed description in [8, 60]) using LEP-MAP3 [63], and a linkage disequilibrium (LD) network based dimensionality reduction (LDna; [64]) was applied to decrease redundancy in the data due to linkage. For each LD-cluster comprising sets of highly correlated SNPs genetic information was extracted using PCA and the PC-coordinates from the first axes explaining the largest proportion of variation were used for QTL mapping. For each cross, we applied the model [8] separately to the phenotypic vectors of size at each age and using the complexity reduced SNP panel after correcting for the effect of sex. We also applied model (8) to the two parameters *k* and *Linf* estimated from the Von Bertalanffy growth curve model described in equation (1). This allowed us to test whether or not there is any global QTL associated with growth in multiple time points.

For each QTL model, a permutation procedure (10000 permutations) was used to account for multiple testing [59, 65].

## Results

### Phenotypic variation in growth trajectories

The asymptotic sizes estimated from the Von Bertalanffy growth curves were similar to the mean body size at the oldest age in HEL x BYN and HEL x PYÖ crosses (Fig. S1; Table S1), suggesting that the data accounted for most of the growth in the measured individuals. In contrast, asymptotic size was higher than mean size at the end of the experiment in HEL x RYT (*Linf* = 55.657 [54.889 - 56.509]; *Lmax* = 48.589 [48.088-49.090]; Table S1). Results from the IDM indicated that variation in ontogeny of body size in all crosses was mainly explained by a growth trajectory predicting an increase in size with age and accounting for 65 - 80 % of the total variation (Fig. S1). In all crosses, a second growth trajectory described a negative correlation between early and late growth (Fig. S1) indicating that some individuals that were larger at an early age tended to be smaller later in life, and *vice versa*.

### Quantitative genetic analyses: genetic variance and heritability

In all crosses and at all ages through ontogeny, there was additive genetic variance and heritability in body size (Fig. 1 & Fig. S2-S4). Additive genetic variance increased with age in all crosses and *V_A_* for body size was significantly higher at the last ontogenetic time-point compared to the first in HEL x PYÖ and HEL x RYT crosses (Fig. 1; Fig. S2-S4; Table S2). SNP-heritability of body size remained relatively constant throughout ontogeny in all crosses and did not significantly differ between ages (Fig. 1; Fig. S2-S4; Table S2).

**Figure 1.**
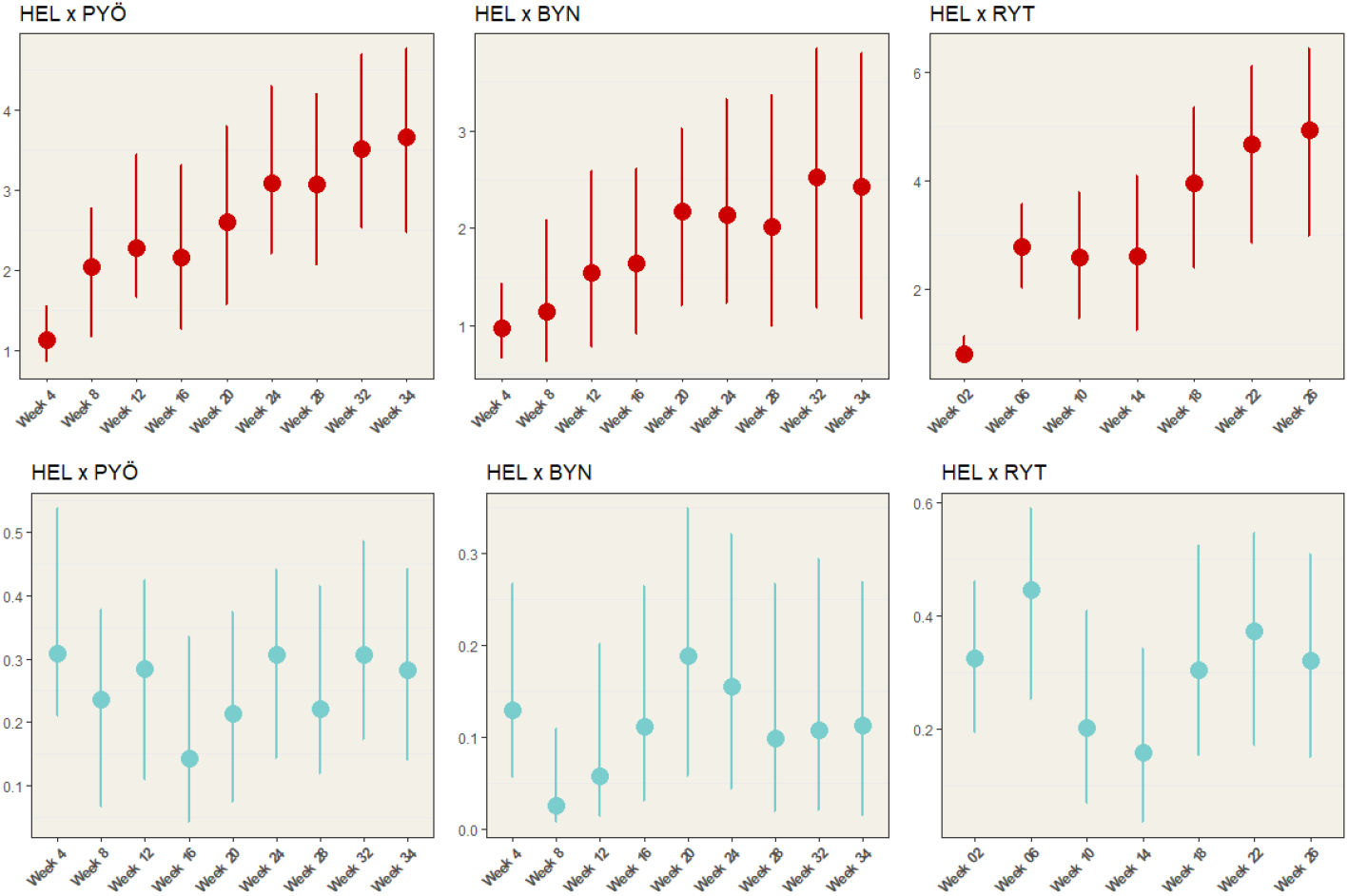
Coefficient of additive genetic variance and heritability for age-specific body size. Posterior modes and 95% HPD intervals for the coefficient of additive variance (*CV_A_*, red) and heritability (*h^2^*, blue) estimated from the *MCMCglmm* animal models are shown for each age-specific body size.

### Quantitative genetic analyses: genetic correlations and G matrix

Genetic correlations between sizes at consecutive ages were positive in all crosses: size at age *n* was positively correlated to size at age *n*+1 and *n*-1 (Fig. 2). Genetic correlations decreased with increasing distance between time-points in all crosses (Fig. 2). In HEL x PYÖ, genetic correlations were no longer statistically different from zero between the size at Week 4 and size at Week 20 and onwards (Fig. 2). In HEL x BYN, body size at first time-point was not genetically correlated to body size at the following time-points (Fig 2). In HEL x RYT genetic correlations decreased with age and body size at the first time-point was not correlated with body size at Week 14, and weakly correlated with body size at Week 18 to 26 (Fig. 2).

**Figure 2.**
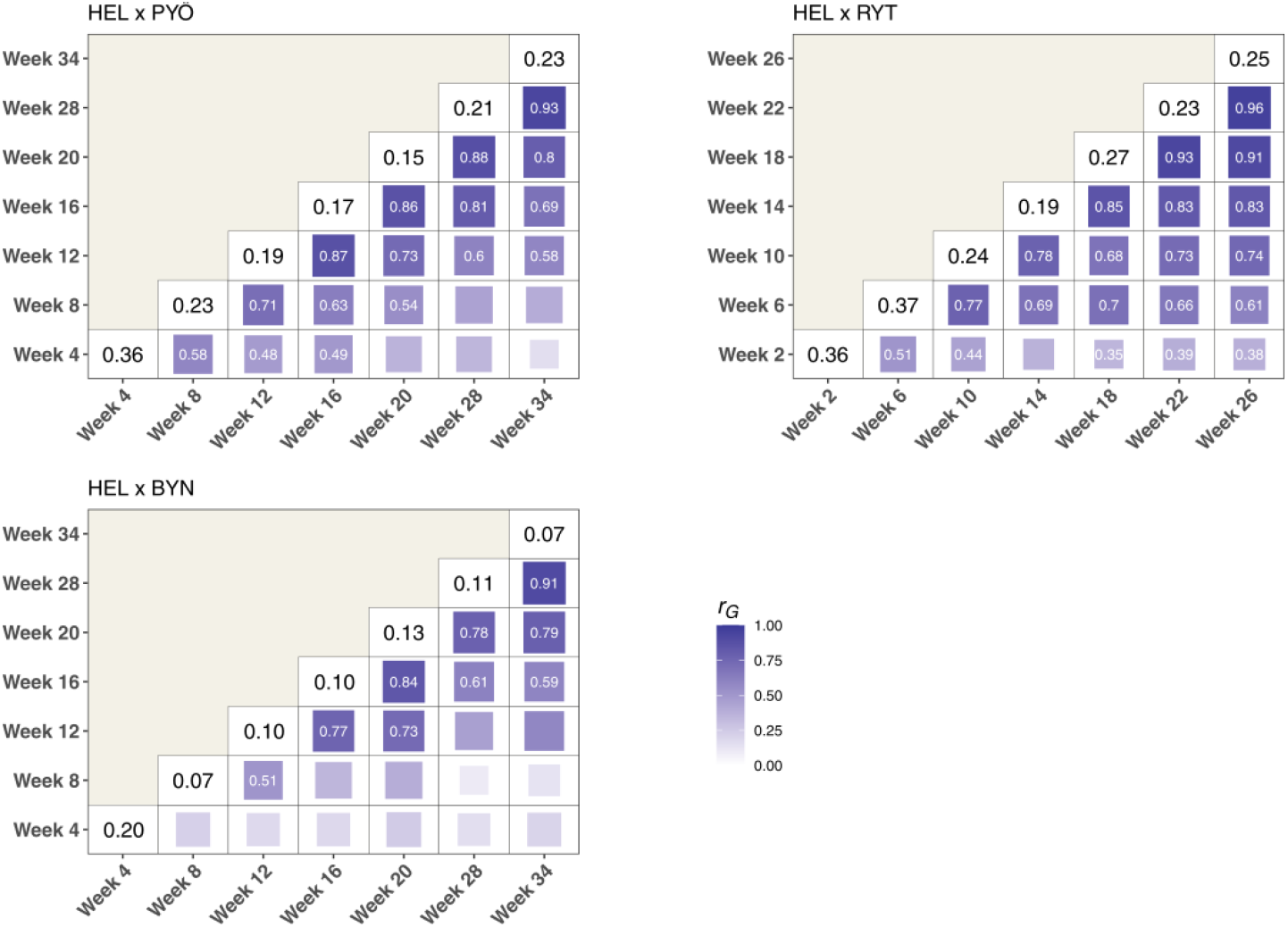
Heritability and genetic correlations across ages. Strength of the genetic correlations (*r_G_*) of body size across ages is represented by the size and colour of each tile of the heatmap. The value of *r_G_* is shown only for significant correlations (white text). The posterior mode of the heritability for body size estimated from the multi-trait model (see *Methods* and also Fig. S3) at each age is shown in the diagonal for each cross (black text).

In all crosses, the first principal component of the **G** matrix explained most of the genetic variance in growth (Table 1), further confirming ample additive genetic variation for body size throughout ontogeny in all crosses. In HEL x PYÖ, a sign change in the loading coefficients of **g**_max_ indicate that early (Week 04, Week 08) and later (Week 16 onwards) sizes are not genetically correlated and that ontogeny is modular between early and late growth. In HEL x BYN, the same result was observed with a sign change of the loading coefficients of **g**_max_ at Week 20. In HEL x RYT loadings were positive for all ages-specific sizes, indicating positive genetic covariation between sizes throughout ontogeny. In HEL x PYÖ and HEL x BYN, a substantial amount of variance (9.73% and 12.36% respectively; Table 1) was also explained by the PC2 where a sign change of the loading coefficients was also observed.

**Table 1.** Percentage of variance explained and loading coefficients of the two first principal components of G. The percentage of genetic variance in growth trajectories explained by the first two principal components of the **G** matrix is shown for each cross. Loading coefficients of **g**_max_ are shown for each age specific body size.

|  | **HEL x PYÖ** |  | **HEL x RYT** |  | **HEL x BYN** |  |
| --- | --- | --- | --- | --- | --- | --- |
|  | PC1 (**g<sub>max</sub>**) | PC2 | PC1 (**g<sub>max</sub>**) | PC2 | PC1 (**g<sub>max</sub>**) | PC2 |
| **% var. explained** | 87.37% | 9.73% | 95.48% | 2.72% | 81.76 % | 12.36% |
| **Age 1** | -0.065 | 0.050 | 0.015 | -0.015 | -0.097 | 0.200 |
| **Age 2** | -0.036 | 0.471 | 0.134 | -0.659 | -0.191 | -0.340 |
| **Age 3** | 0.137 | 0.621 | 0.185 | -0.680 | 0.074 | -0.627 |
| **Age 4** | 0.263 | 0.480 | 0.295 | -0.153 | 0.260 | -0.489 |
| **Age 5** | 0.415 | 0.198 | 0.450 | 0.153 | 0.406 | -0.321 |
| **Age 6** | 0.556 | -0.099 | 0.547 | 0.168 | 0.570 | 0.113 |
| **Age 7** | 0.652 | -0.334 | 0.599 | 0.166 | 0.626 | 0.309 |

### QTL-mapping

A total of 11 unique QTL associated with body size variation at different ontogenetic stages were detected (Fig. 3). In the HEL x BYN cross three different QTL on chromosome (Chr.) 9 (Week 16 to 28), Chr. 10 (Week 12, 16) and Chr. 18 (Week 12) were found (Fig. 3A). In HEL x PYÖ six QTL on Chr. 1 (Week 34), Chr. 9 (Week 4 and Week 12 to 28), Chr. 13 (Week 4), Chr. 14 (Week 4), Chr. 18 (Week 12) and Chr. 20 (Week 32 & 34) were discovered (Fig. 3B). In HEL x RYT four QTL localized in Chr. 8 (Week 6), Chr. 12 (Week 2 & 6), Chr. 15 (Week 6) and Chr. 17 (Week 6 & 10) were significant (Fig. 3C). We did not find any global QTL underlying the growth parameters *k* and *Linf* in the HEL x BYN cross (Fig. S5). A single QTL was detected in HEL x PYÖ for *k* on Chr. 1 (Fig. S5) and similarly for HEL x RYT, a single QTL controlling for asymptotic size (*Linf*) was found on Chr. 7 (Fig. S5).

**Figure 3.**
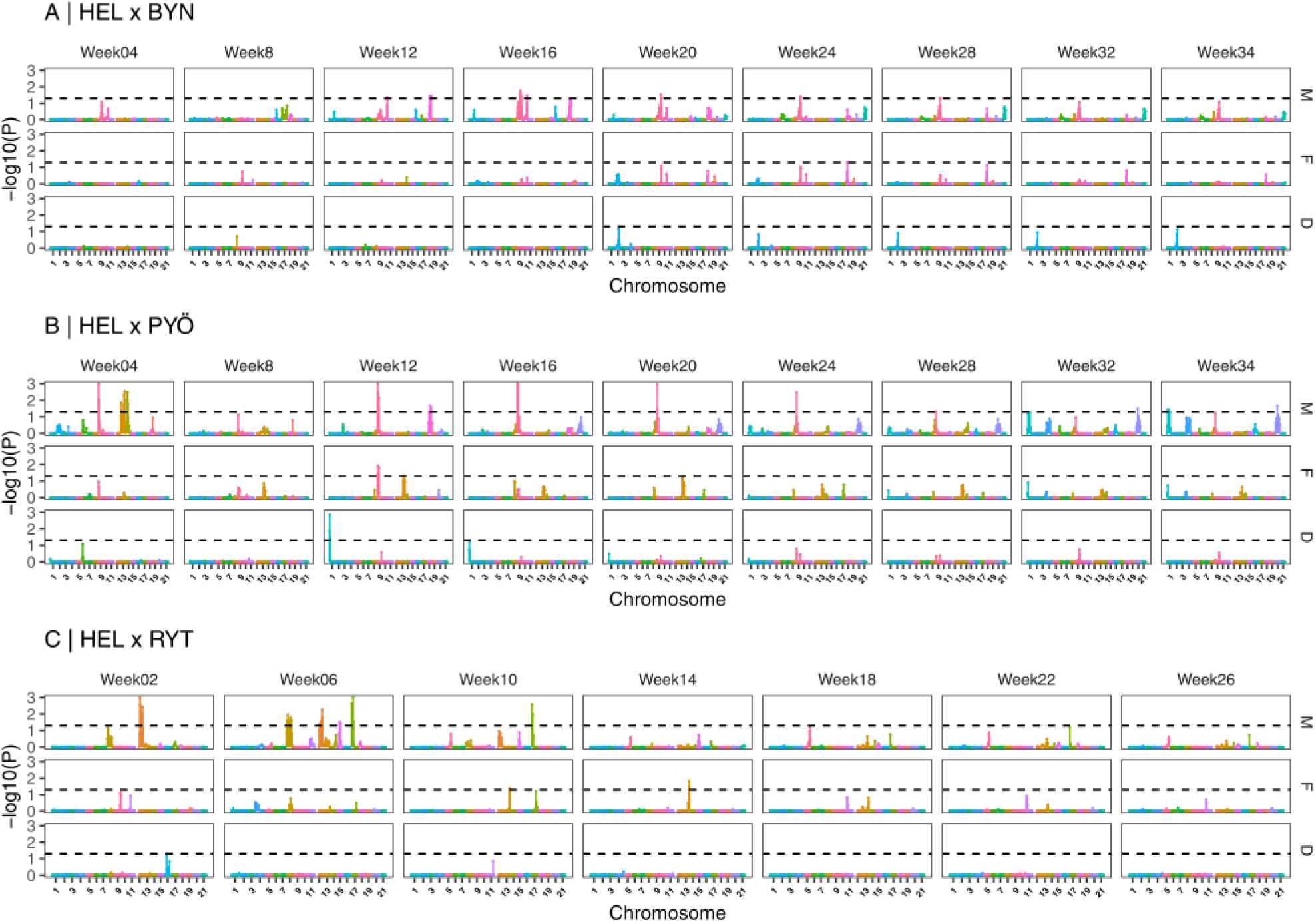
QTL-mapping results for age-specific body size. Results from the four-way QTL-mapping are shown for each cross (A,B,C) and for body size at each age (Week *n*). For each cross and each age, panels show whether the QTL are inherited from the sire (M) or the dam (F) along with the dominance effect (D) estimated from model (8) in the main text. Results are based on permutation and the significance threshold (dashed horizontal line) is shown on the logarithm scale (*p* = 0.05). Colors represent different chromosomes.

This shows that different QTL underline expression of body size not only among the different crosses but also between different ontogenetic changes within the crosses (Fig. 3). Most of the QTL effects traced down to the sire (M, in Fig. 3) indicating that the observed allelic effects were segregating in the male F_0_ grandfather (pond individual). One peak indicative of segregation in the F_0_ female (marine individual) on Chr. 9 (at age Week 12 in HEL x PYÖ; Fig. 3B), and another on Chr. 13 (at ages Week 10 and 14 in the HEL x RYT cross; Fig. 3C) were found in addition to the QTL for *k* and *Linf* in HEL x PYÖ and HEL x RYT, respectively. Dominance effects were rare: only a single significant dominance allelic effect on Chr. 1 at age Week 12 in the HEL x PYÖ cross was observed (Fig. 3B).

## Discussion

Heterogeneous genetic underpinnings for similar phenotypic adaptations are a testimony of redundancy of organismal genetic architectures: similar phenotypic end-points can be reached with diverse genetic mechanisms. Our results demonstrate that in spite of very similar phenotypic growth trajectories and approximately constant heritabilites of body size across different ages and crosses, the QTL contributing to size were different across ages both within and among the three crosses studied here. These results were supported by analyses of genetic correlations across the age groups; early and late growth genetic correlations were generally weak or not significantly different from zero, and always significantly lower than one. These results align with earlier findings from this study system showing that heterogeneous genetic architectures underlie similar phenotypic adaptations in different pond populations [8]. The main novelty in the current results is that they demonstrate ontogenetic heterogeneity in genetic architecture of an important life history trait; would selection act on size at different points of development, genes governing the expected selection responses would be very different. In the following, we will discuss these and related points in more detail.

Within each of the three crosses different QTL were often associated with body size at different ages, and the significant QTL effects were predominantly associated with alleles derived from the pond grandparent, and seldom with alleles from the marine grandparent. The latter finding makes sense in the light of the fact that pond fish have been shown to be consistently larger than marine fish throughout development [39], and therefore, one would expect pond alleles to contribute to size more strongly than the marine alleles. In few cases QTL effects tracing to both parents were observed, and only in one case was there evidence for dominance contribution. Hence, these findings suggest that most observed allelic effects were additive, and mostly coming from pond genetic background.

The ontogenetic heterogeneity in the QTL effects was clear in all three crosses and the results of the analyses of genetic correlations supported the inference; different genetic architectures underline expression of body size variation in different ontogenetic time-points and no global QTL seemed to govern the expression of size across these points. We observed also among cross heterogeneity in the age-specific QTL effects, suggesting that genetic architecture of body size variability might differ among the pond populations similarly as shown earlier for pelvis reduction in these populations [8]. However, some caution is warranted in interpretation of the observed population differences in QTL locations for two reasons. First, while exact time-points of measurement were the same for two of the three crosses, fish in one of the crosses were measured in slightly different time-points than in the two others. Given that we picked up time-dependent variation in QTL effects, this could have influenced which QTL were detected in one of the crosses. However, the similarly measured crosses showed different patterns of QTL expression. Second, it should be kept in mind that all three crosses were established from a single pond and a single marine (grand) parent, and the crosses are therefore unlikely to carry all relevant allelic variation present in the populations of origin. Hence, there might be a large stochastic component to QTL detectable in each of the three crosses. While we do not have data to refute this possibility, an earlier study using empirical and simulated data gave strong evidence for heterogeneous genetic architecture underlying pelvic reduction in these populations [8, 66], thus suggesting that this stochasticity might not be of great concern. Irrespectively of what explains heterogeneous QTL effects among different crosses, sampling of allelic variation in parental populations is irrelevant to interpretation of the heterogeneous QTL effects across ontogeny.

We found additive genetic variance and heritability for body size across ages in all three crosses. This suggests that body size can respond to directional selection at different stages of the ontogeny. However, genetic correlations between age-specific sizes tended to decay with time, and the correlations between early and late sizes were weak or non-existent. This was also manifested in the analyses of integration of the growth trajectory, which revealed evidence for modular genetic architecture of growth in two of the three crosses. This means that selection on size at any given age would not necessarily affect size across the whole ontogeny [49, 58]. On the contrary, the results suggest that genes responsible for early and late growth are acting independently, and therefore, selection acting at early and later ages would be acting on different sets of genes. These results align with the results of QTL analyses which show that different QTL control body size in different ages, and there were no global QTL which would influence growth (as reflected in *k*) in two of the three crosses.

The variance component and heritability estimates in this study were based on single full-sib family per cross utilising variance in relatedness among sibs to estimate the causal components of variance. This approach has been utilized in earlier studies (*e.g*. [42, 67]) and has the advantage that the estimates are not subject to additional variance attributable to uncontrolled environmental and maternal effect sources. That said, since the analysed crosses are artificial F_2_ generation inter-population crosses, the estimated quantitative genetic parameters may not be representative about those in wild populations. Regardless, the main results - approximately constant heritability but varying genetic architecture over the ontogeny - are indisputable facts providing a conceptual framework to think how ontogenetic changes in genetic architecture may influence predictability of evolution in nature.

In conclusion, using both quantitative genetic and QTL-mapping tools, we have shown that while the heritability of body size remains relatively constant across the ontogeny, its underlying genetic architecture does not. While there are many examples of heterogeneous genetic architectures underlining similar phenotypic expression of traits (*e.g*. [8, 66, 68]), few studies from the wild have been able to demonstrate this for the same trait across ontogenetic stages. This ontogenic genetic heterogeneity suggests that if directional natural selection would act on body size at different ages in two different populations, the outcome would be divergence in genetic architecture of body size in these populations. Hence, ontogenetic differences in genetic architecture underlying expression of genetic variation are a potential source of evolution of heterogeneous genetic architecture among different populations of the same species.

## Acknowledgments

Thanks are due to Gabor Herczeg, Abigel Gonda, Yukinori Shimada, Mirva Turtiainen, Chris Eberlein, Takahito Shikano, Laura Hänninen, Kirsi Kähkönen, Miinastiina Issakainen and Sami Karja for fish breeding and DNA extractions and Jing Yang for help in phenotyping the fish. The authors thank Petri Kemppainen and Emma Vatka for useful discussions on previous versions of the manuscript. The computing resource support from CSC - the Finnish IT Center for Science Ltd administered by the Ministry of Education and Culture, Finland is gratefully acknowledged.

## Ethics

All experimental protocols were approved by permission (ESLHSTSTH223A) form the National Animal Experiment Board, Finland.

## Author contribution

J.M. and A.F. conceived the study; Z.L. contributed to pre-processing and getting the genotype data; Z.L and M.S. provided analytical resources; A.F. analyzed the data; J.M. and A.F. led the writing of the manuscript. All authors contributed critically to the drafts and gave final approval for publication.

## Data availability

Raw sequence reads have been submitted to NCBIs short-read archives with accession nos.: PRJNA673430 and PRJNA672863. Phenotype data will be deposited to Dryad upon acceptance.

