## Supplementary material for "Age-dependent genetic architecture underlines similar heritability of body size in sticklebacks"

Running head: Age dependent genetic architecture in sticklebacks

**Table of contents**


| | |
|---|---|
| **Figure S1**. Phenotypic variation in growth curves | page 2 |
| **Figure S2**. Additive genetic variance and heritability estimated by Gaussian Process | page 3 |
| **Figure S3**. Additive genetic variance and heritability estimated from the multi-trait model | page 4 |
| **Figure S4**. Penalized spline smoothing of additive genetic variance and heritability | page 5 |
| **Figure S5**. QTL-mapping results for growth rate (*k*) and asymptotic size (*Linf*) | page 6 |
| **Table S1**. Von Bertalanffy growth parameters for the three crosses | page 7 |
| **Table S2**. Quantitative genetic parameters of age-specific body size | page 8 |

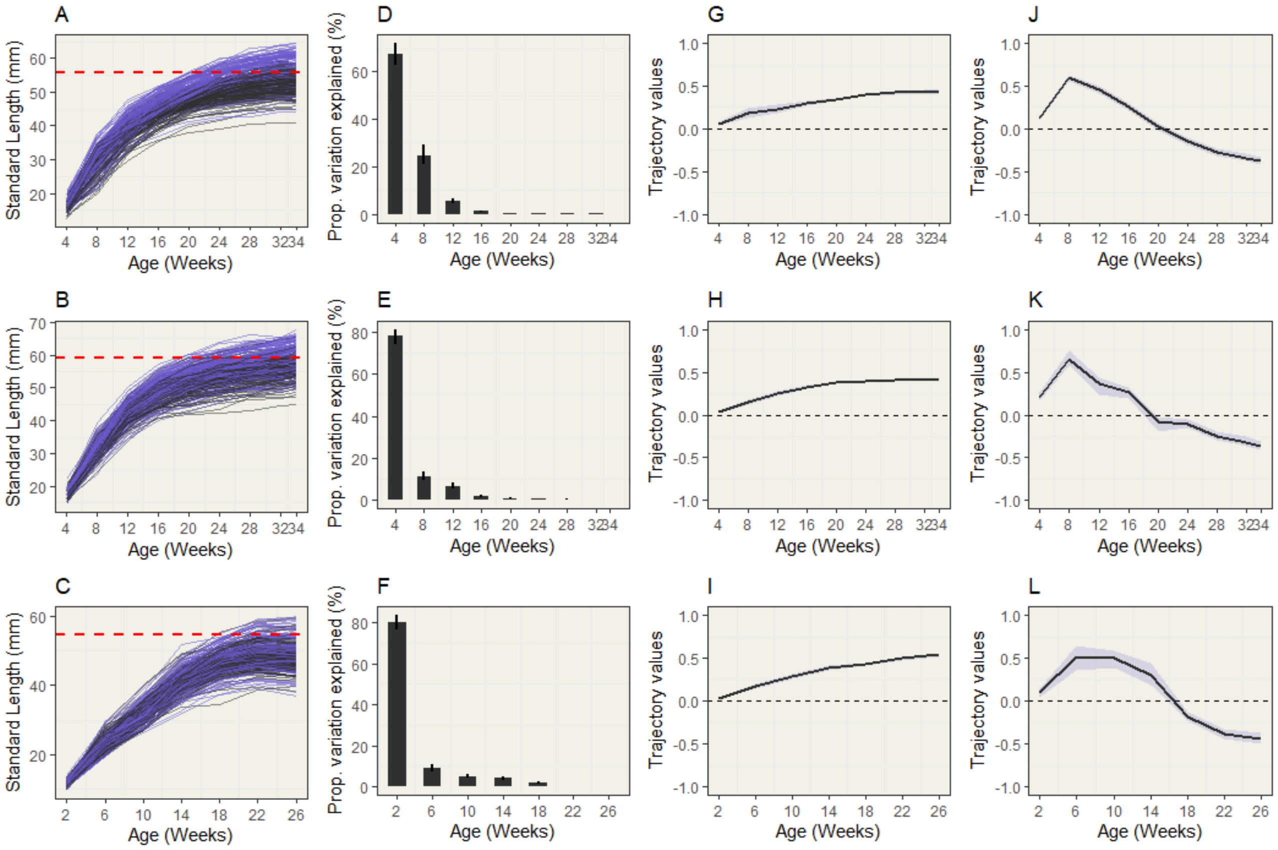


**Figure S1. Phenotypic variation in growth curves.** Individual growth trajectories are shown for males (black solid lines) and females (purple solid lines) of the HEL x BYN (A), HEL x PYÖ (B) and HEL x RYT (C) crosses along with the asymptotic size (red dashed line) estimated from the VBGC model (equation (1) in main text). IDM applied to HEL x BYN (D, G, J), HEL x PYÖ (E, H, K) and HEL x RYT (F, I, L) growth data. The percentages of phenotypic variation (95% CIs encompassed by vertical lines) accounted for by each growth trajectories is shown by the barplots. The first (G, H, I) and second (J, K, L) trajectory are shown for each cross (black solid lines) along with 95% CIs for the trajectories (purple shadings).

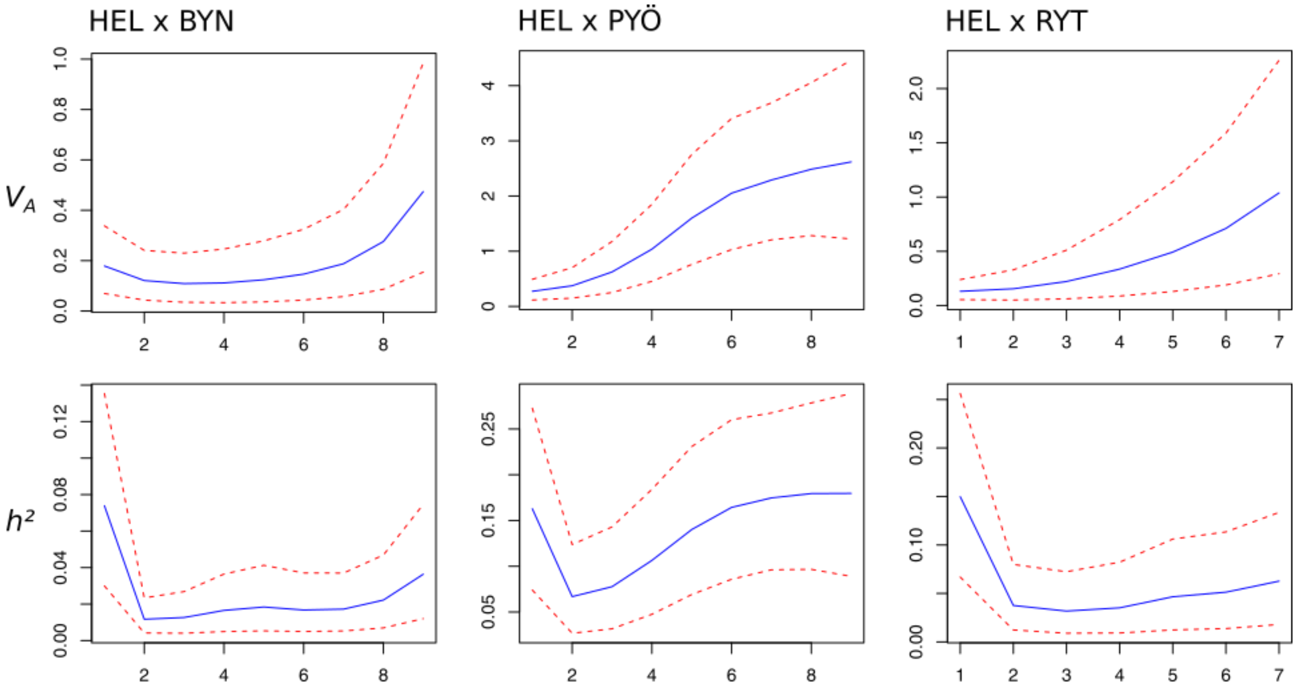


**Figure S2. Additive genetic variance and heritability estimated by Gaussian Process.** Posterior mean curves (blue, solid lines) and 95% credible intervals (red, dashed line) are drawn for the estimation of genetic variance ($V_A$, left panels) and heritability ($h^2$, right panels) for each cross.

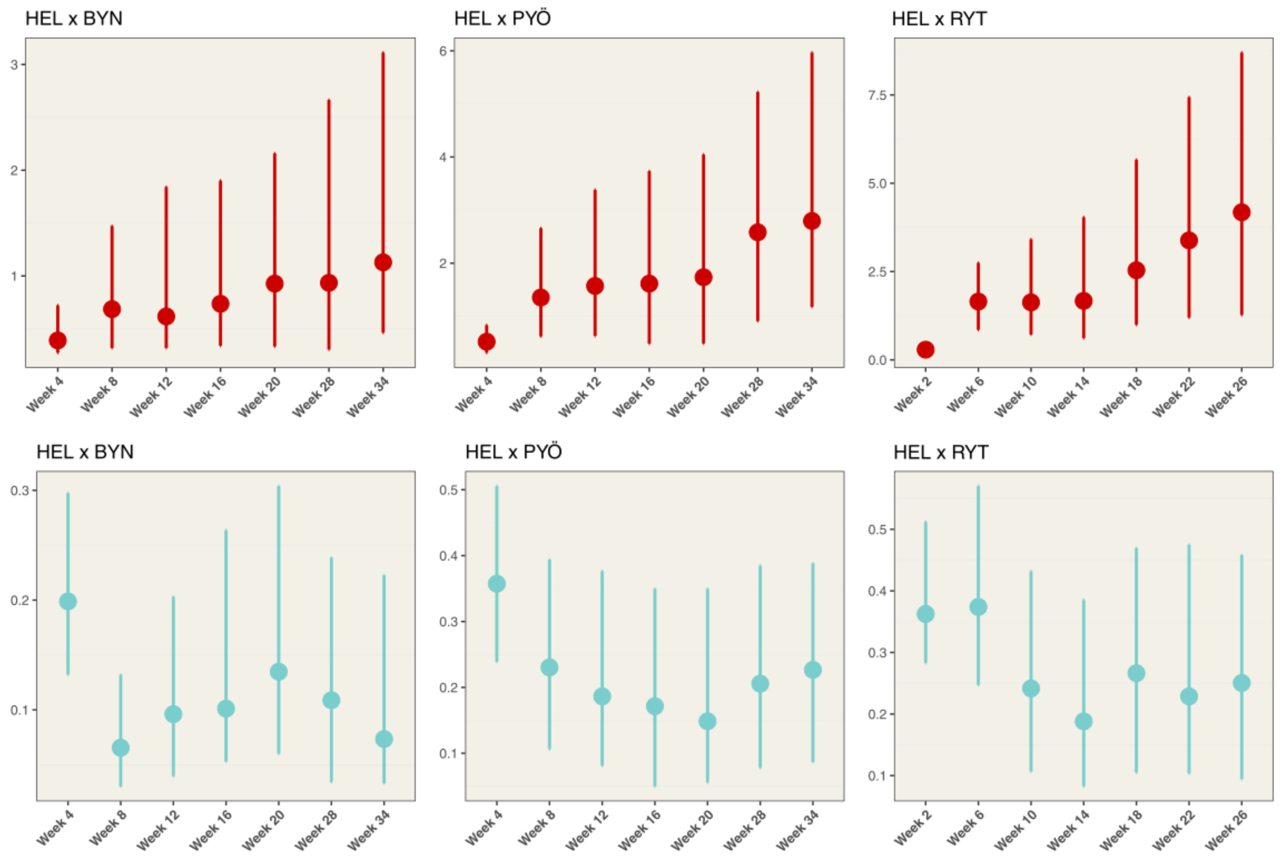


**Figure S3. Additive genetic variance and heritability estimated from the multi-trait model.** Posterior modes and 95% HPD intervals for additive variance ($V_A$, red) and heritability ($h^2$, blue) estimated from the *MCMCglmm* multivariate animal models (see *Methods*) are shown for each age-specific body size.

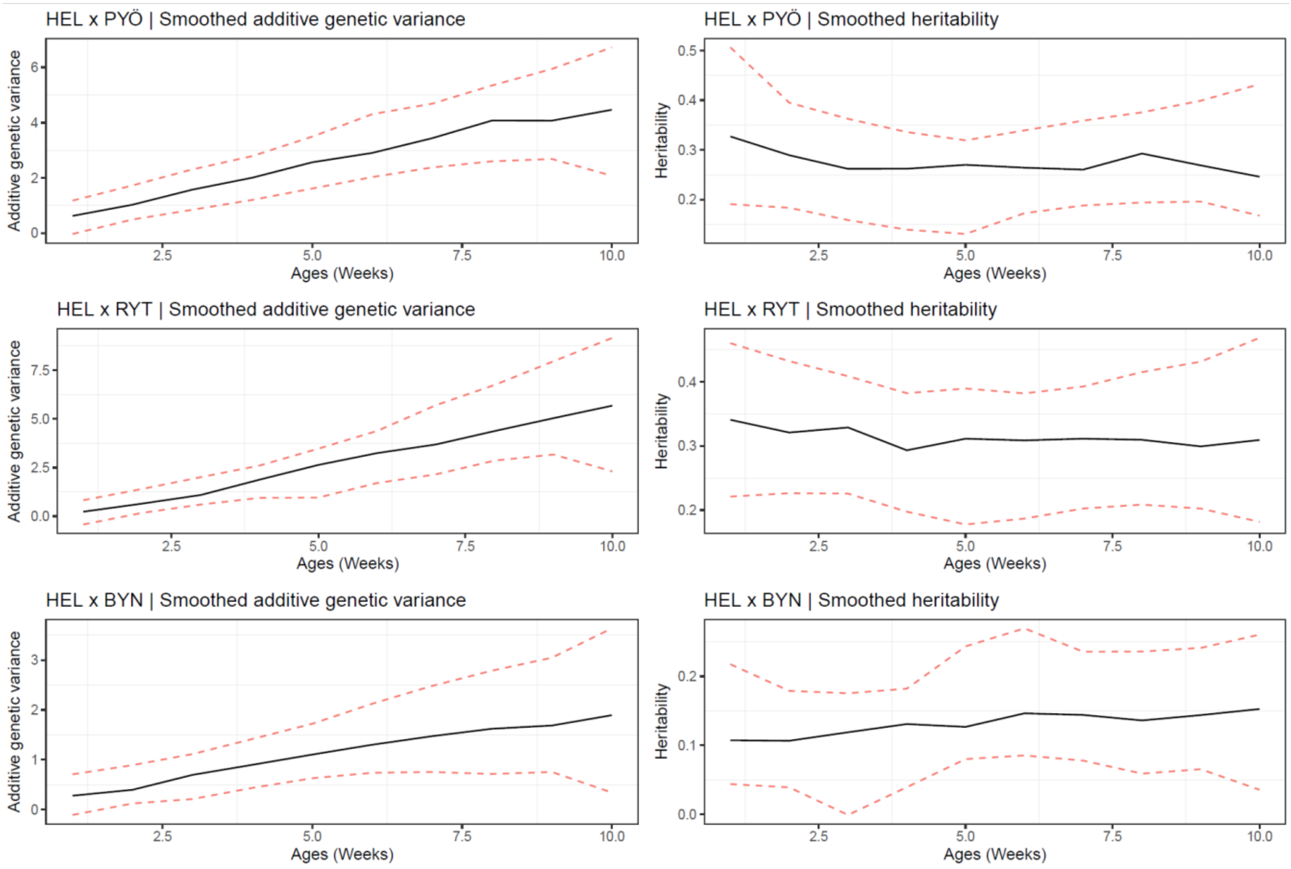


**Figure S4. Penalized spline smoothing of additive genetic variance and heritability.** Posterior modes (black solid lines) and 95% HPD intervals (red dashed lined) are shown for smoothed values of additive variance ($V_A$) and heritability ($h^2$) estimated from the *MCMCglmm* univariate animal models (see *Methods*). Smoothed values are plotted along the 10 basis functions of the penalized spline function corresponding to the time points along development.

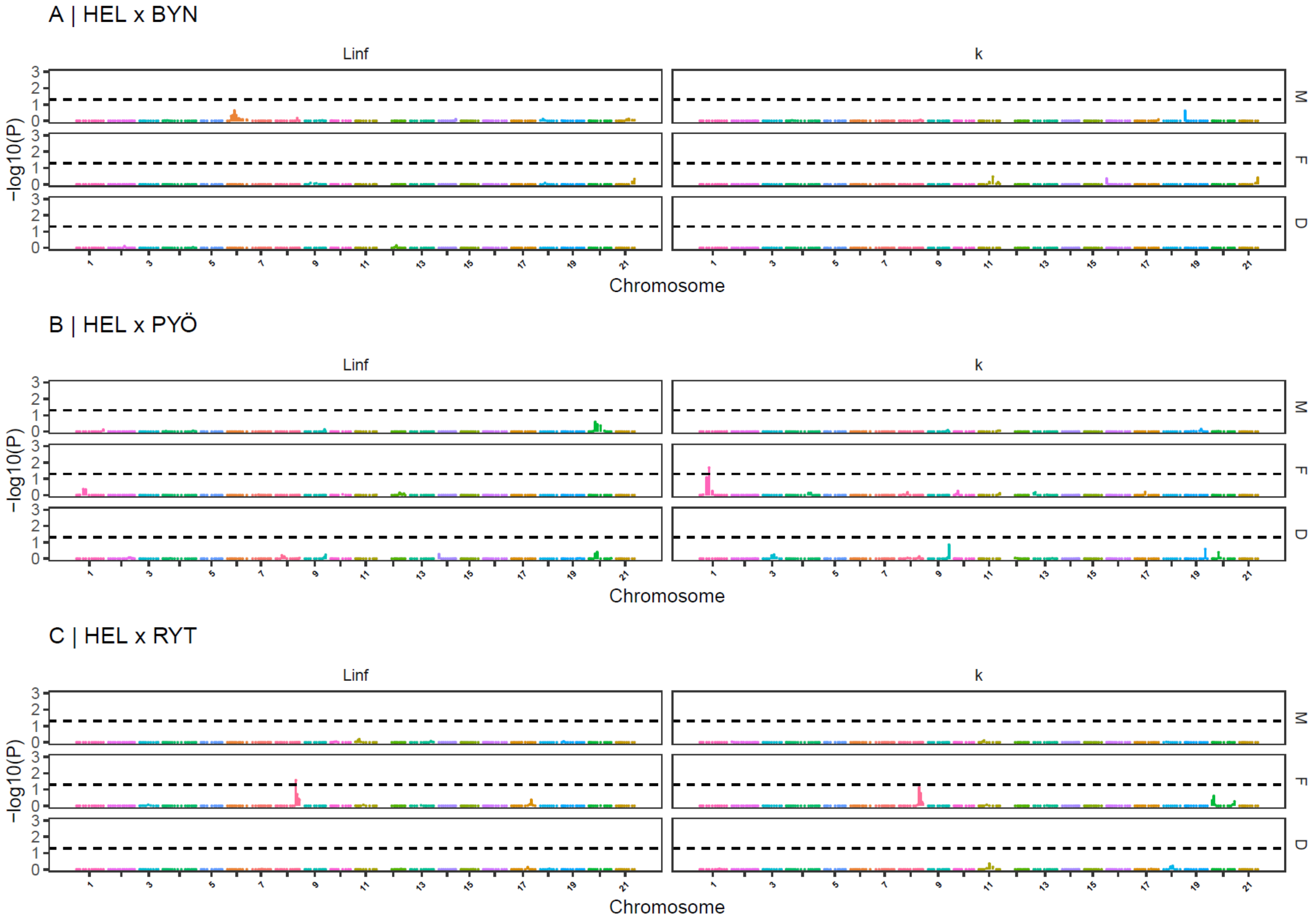


**Figure S5. QTL-mapping results for growth rate (*k*) and asymptotic size (*Linf*).** Results from the four-way QTL-mapping are shown for each cross (A,B,C) and for parameters *k* and *Linf* estimated from equation [1] in the main text. For each cross and each trait, panels show whether the QTL are inherited from the sire (M) or the dam (F) along with the dominance effect (D) estimated from model (8) in the main text. Results are based on permutation and the significance threshold (dashed horizontal line) is shown on the logarithm scale ($p$ = 0.05). Colors represent different chromosomes

**Table S1. Von Bertalanffy growth parameters for the three crosses.** Growth parameters and their 95% confidence intervals estimated from bootstrap are shown for the growth rate *k* and the asymptotic length $L_{inf}$ as in equation (1) in the main text. $L_{max}$ corresponds to the mean body size measured at the last time point of the experiment (Week 34 for HEL x BYN and HEL x PYÖ; Week 26 for HEL x RYT) and the 95% CI estimated from a one sample *t-test*.

| | *k* | $L_{inf}$ | $L_{max}$ |
|---|---|---|---|
| HEL x BYN | 0.111 [0.108 - 0.114] | 55.768 [55.440 - 56.091] | 54.162 [53.693-54.630] |
| HEL x PYÖ | 0.115 [0.112 - 0.118] | 58.863 [58.545 - 59.180] | 57.782 [57.302-58.263] |
| HEL x RYT | 0.089 [0.085 - 0.093] | 55.657 [54.889 - 56.509] | 48.589 [48.088-49.090] |

**Table S2. Quantitative genetic parameters of age-specific body size.** Posterior modes and 95% HPD intervals are shown for the variance components estimated from the univariate animal models (equation (2) in *Methods*) using *MCMCglmm*: additive genetic variance ($V_A$), residual variance ($V_R$), phenotypic variance ($V_P$) and heritability ($h^2$) are given for each age in each cross.

| Cross | Age | $V_A$ | $V_R$ | $V_P$ | $h^2$ |
|---|---|---|---|---|---|
| HEL x BYN | Week 04 | 0.279 [0.113; 0.582] | 1.690 [1.417; 2.051] | 2.005 [1.731; 2.401] | 0.129 [0.056; 0.266] |
| | Week 08 | 0.266 [0.086; 1.215] | 10.209 [8.802; 12.226] | 10.621 [9.275; 12.728] | 0.025 [0.008; 0.108] |
| | Week 12 | 0.464 [0.113; 1.826] | 7.637 [6.401; 9.249] | 8.706 [7.349; 10.141] | 0.057 [0.014; 0.201] |
| | Week 16 | 0.792 [0.153; 1.834] | 5.674 [4.691; 6.909] | 6.703 [5.640; 7.830] | 0.111 [0.031; 0.264] |
| | Week 20 | 1.173 [0.309; 2.487] | 5.274 [4.254; 6.424] | 6.594 [5.560; 7.867] | 0.189 [0.057; 0.349] |
| | Week 24 | 1.343 [0.339; 3.031] | 7.400 [5.855; 8.758] | 8.676 [7.418; 10.391] | 0.155 [0.043; 0.321] |
| | Week 28 | 1.189 [0.139; 3.072] | 9.614 [7.683; 11.397] | 10.659 [9.289; 12.913] | 0.099 [0.018; 0.267] |
| | Week 32 | 1.402 [0.245; 3.987] | 10.308 [8.436; 12.714] | 12.160 [10.482; 14.678] | 0.107 [0.020; 0.294] |
| | Week 34 | 1.374 [0.180; 3.857] | 11.309 [9.375; 14.005] | 13.390 [11.364; 15.788] | 0.113 [0.014; 0.270] |
| HEL x PYÖ | Week 04 | 0.413 [0.225; 0.791] | 0.799 [0.606; 0.972] | 1.163 [1.027; 1.554] | 0.308 [0.209: 0.538] |
| | Week 08 | 1.034 [0.452; 2.591] | 4.569 [3.635; 5.516] | 5.758 [4.895; 7.01] | 0.236 [0.066; 0.378] |
| | Week 12 | 1.736 [0.826; 3.815] | 5.998 [4.824; 7.37] | 8.035 [6.722; 9.732] | 0.284 [0.110; 0.424] |
| | Week 16 | 1.573 [0.373; 3.421] | 7.609 [6.164; 9.390] | 9.443 [8.065; 11.45] | 0.142 [0.042; 0.334] |
| | Week 20 | 2.271 [0.700; 4.611] | 8.461 [7.069; 10.709] | 11.39 [9.327; 13.641] | 0.213 [0.074; 0.374] |
| | Week 24 | 3.203 [1.528; 6.019] | 8.727 [6.741; 10.526] | 12.39 [10.191; 14.602] | 0.306 [0.143; 0.441] |
| | Week 28 | 3.159 [1.202; 5.654] | 8.851 [7.254; 11.063] | 12.874 [10.284; 14.83] | 0.220 [0.119; 0.415] |
| | Week 32 | 3.186 [1.943; 7.071] | 9.165 [7.06; 10.938] | 14.014 [11.041; 15.979] | 0.306 [0.173. 0.486] |
| | Week 34 | 4.474 [1.715; 7.030] | 9.927 [7.941; 12.230] | 14.178 [11.863; 17.242] | 0.282 [0.139; 0.443] |

| | | | | | |
|---|---|---|---|---|---|
| HEL x RYT | Week 02 | 0.161 [0.095; 0.293] | 0.379 [0.304; 0.477] | 0.575 [0.484; 0.707] | 0.326 [0.193; 0.460] |
| | Week 06 | 1.835 [0.836; 2.855] | 2.397 [1.836; 3.042] | 4.161 [3.452; 5.249] | 0.446 [0.252; 0.591] |
| | Week 10 | 1.605 [0.364; 3.170] | 4.996 [4.183; 6.538] | 6.680 [5.794; 8.431] | 0.202 [0.068. 0.409] |
| | Week 14 | 1.604 [0.258. 3.596] | 7.956 [6.657; 10.157] | 9.592 [8.282; 11.895] | 0.157 [0.035. 0.341] |
| | Week 18 | 3.691 [1.150; 6.527] | 6.793 [5.475; 9.106] | 11.159 [9.101; 13.641] | 0.304 [0.152. 0.526] |
| | Week 22 | 4.465 [1.743; 8.560] | 9.463 [7.496; 12.288] | 14.271 [11.799; 17.758] | 0.374 [0.171; 0.546] |
| | Week 26 | 5.737 [1.888; 9.478] | 11.625 [9.376; 15.222] | 17.356 [14.707; 21.565] | 0.321 [0.148; 0.509] |
